# Temperature modulates the dissemination potential of *Microsporidia MB,* a malaria-blocking endosymbiont of *Anopheles* mosquitoes

**DOI:** 10.1101/2024.11.15.623820

**Authors:** Fidel Gabriel Otieno, Priscille Barreaux, Affognon Steeven Belvinos, Edward Edmond Makhulu, Thomas Ogao Onchuru, Anne Wambui Wairimu, Stancy Mandere Omboye, Cynthia Nyambura King’ori, Sokame Bonoukpoè Mawuko, Anthony Nyamache Kebira, Jeremy Keith Herren

## Abstract

The endosymbiont *Microsporidia MB* is a promising malaria control strategy that inhibits the development of *Plasmodium* naturally in *Anopheles* mosquitoes. To be successful, it would be necessary to significantly increase the prevalence of *Microsporidia MB* in populations of malaria mosquitoes to decrease the malaria transmission potential of the mosquito population. However, very little is known about the role of temperature in driving the prevalence of *Microsporidia MB* infections in mosquito populations. By rearing mosquito larvae under four air temperature regimes (22°C, 27°C, 32°C and 37°C), we show that warm temperatures favour the growth of *Microsporidia MB* infected larvae. In addition, *Microsporidia MB* infected larvae developed faster compared to the uninfected offspring of the same mothers. Starting with 10 *Microsporidia MB* infected mothers, our population growth model showed that, at 32°C, it would take 15-35 days to reach a population of 1000 *Microsporidia MB* infected mothers; this represents a dissemination potential of 4.7, 1.3 and 1.7 times higher compared to 22°C, 27°C and 37°C, respectively. Despite a relatively high mosquito mortality rate (20% more compared to 27°C), 32°C was estimated the best temperature for rearing *Microsporidia MB* infected larvae due to the shorter development time and high infection rate. This study gives insight into the favourable conditions for *Microsporidia MB* mass rearing and potential release strategies in malarious regions.

**Importance:** Malaria parasites transmitted by *Anopheles* mosquitoes cause a life-threatening disease, imposing a massive toll on human health and economic sustainability in sub-Saharan Africa. Relying only on insecticide- and drug-based control products whose efficacy has been eroded by resistance to control malaria is not sufficient anymore. New innovative approaches are urgently needed and *Microsporidia MB*, a naturally occurring symbiont across Africa is capable of inhibiting *Plasmodium* transmission in *Anopheles gambiae* s.l.. Its success in adverting a rebound of malaria cases will depend on the infection dynamic of the symbiont over time and space. Through experimental studies on field derived mosquitoes and mathematical modelling, we demonstrate that *Microsporidia MB* dissemination potential increase with temperature within a viable range for *Anopheles* mosquitoes, due to trade-offs between mosquito development and survival and the symbiont growth. Future studies should now investigate how fluctuating temperatures modulate the *Plasmodium* transmission blocking performance in nature.

## Introduction

Malaria prevention management is impacted by increasing temperature globally (1) and the threat of widespread insecticide resistance particularly in many low- and middle-income countries in sub-Saharan Africa (2). While climate warming promotes insecticide resistance and impedes the efficacy of current insecticide-based control measures (3), the number of malaria cases worldwide is alarmingly increasing with an estimated 249 million cases in 2022 compared to 231 million cases in 2015 (4). The critical targets of the World Health Organization (WHO) to achieve 90% reduction of malaria cases incidence and mortality rates by 2030 compared with the 2015 baseline cannot be met with current insecticide-based control tool measures alone (5). Complementary and novel malaria control innovations that address the issue of resistance are needed and their efficacy should be evaluated considering seasonality, the ecological profile of the vectors and climate-dependant tolerance to insecticide (4).

*Microsporidia MB* is a promising novel malaria control strategy that naturally inhibits *Plasmodium* parasite proliferation in *Anopheles* mosquitoes (6, 7) and is spreading in nature and mosquito populations using vertical and horizontal transmission routes (6). While this symbiont is currently under investigation to determine its dissemination and malaria blocking performance in nature, investment is needed to understand the impact of temperature and climate on this promising malaria control strategy. In fact, *Microsporidia MB* shows seasonal variation in prevalence with highest level of infected mosquitoes after peak rainfalls (7), as it is observed in several microsporidians in mosquitoes (8).

In particular, high temperature and low humidity promote the growth of microsporidians by increasing the infection intensity and production of spores (8–11). The physiological state and behaviour of mosquitoes is also dependent on climate and environmental conditions (12–14). Temperature strongly affects mosquito fitness (8,15–18) and influences adult size, survival, breeding, and reproduction (17,18,19–21). In other insect species, temperature has been shown to play an important role in symbiosis. *Nezara viridula*, a stinkbug pest, experiences a fitness cost in warmer conditions, which apparently suppresses its gut microbiome (22). As *Microsporidia MB* transmission is intensity-dependent (6,7) and the intensity of infection can be affected by mosquito age and blood feeding (7,23). Temperature can impact the proliferation of *Microsporidia MB* and hence present opportunities or challenges for the development and implementation of a *Microsporidia MB-*based malaria control approach. In this study, we investigated the impact of temperature on the dissemination potential of *Microsporidia MB*-infected *Anopheles arabiensis* mosquitoes based on the mosquito larval development time and survival as well as *Microsporidia MB* infection rates in mosquito offspring.

A continuous logistic model was chosen to provide a smooth and accurate representation of mosquito population growth, reflecting natural, gradual changes without the constraints of fixed time intervals required by discrete models. This continuous approach offers precise population estimates at any point in time, making it ideal for understanding temporal growth rates and allowing for the straightforward incorporation of stochastic variability to reflect environmental influences on fecundity. A carrying capacity K of 1000 was set to simulate real-world limitations such as resource and space constraints, establishing a stable population maximum that aligns with natural conditions. Additionally, targeting a population of 1000 *Microsporidia MB* positive offspring provides a measurable endpoint for assessing the spread of *Microsporidia MB* within mosquito populations, offering insights into its potential as a vector control tool. By tracking the time required to reach this threshold, our model yields valuable data on the speed and dynamics of *Microsporidia MB* dissemination across generations, enhancing understanding of *Microsporidia MB*’s effectiveness in vector control.

## Results

### 1) Quantification of *Microsporidia MB*

Since the efficiency of *Microsporidia MB* vertical transmission is variable, the progeny of *Microsporidia MB* infected females contains both *Microsporidia MB* positive and *Microsporidia MB* negative mosquitoes. Overall, the vertical transmission rate observed in this study was 43%; *Microsporidia MB* positive and negative progeny from infected mothers have been respectively confirmed positive and negative by PCR screening. Since horizontal transmission of *Microsporidia MB* has so far not been observed during larval development, the offspring of females from *Microsporidia MB* negative mothers are all considered un-infected.

#### Pupation rate

across all experiments the average pupation rate in the offspring was 54.1 (95% confidence interval: 51.60-56.59) %. At 27°C, we observed a pupation rate of 73.8 (69.44-78.14) % which was not significantly different from 22°C; but 18 % and 55 % higher than that observed at 32°C and 37°C, respectively [22°C: 67.8 (63.16-72.36) %; 32°C: 56.1 (50.92-61.13) %; 37°C: 19.0 (15.12-22.86) %, **(Figure 1.1)**] (Tukey post-hoc comparisons: p_27°C-22°C_ > 0.05 and p_27°C-32°C_ and p_27°C-37°C_ < 0.001; *χ*^2^= 153.95, df = 3, p < 0.001). For *Microsporidia MB* positive mothers, the difference between the pupation rates in offspring at different temperatures were all significantly difference from 27°C and presented a positively skewed bell shaped distribution: the pupation rate was best at 27°C, 74.4 (70.14-79.56) %; then 22°C, 68.4 (63.37-73.37) %; then 32°C, 55.8 (49.84-60.95) % and low at 37°C, 14.6 (10.77-18.40) % (Tukey for MB+ mothers: p_22°C-27°C_ = 0.001; p_27°C-32°C_ = 0.137, p_27°C-37°C_ < 0.001; p_32°C-37°C_ < 0.001; p_22°C-32°C_ > 0.05; *χ*^2^= 29.39 df = 3, p < 0.001). The overall better pupation rate in offspring of *Microsporidia MB* negative mothers compared to *Microsporidia MB* positive mothers [MB-mothers: 58.5 (52.29-64.73) %; MB+ mothers: 53.3 (50.56-56.00) %] was only due to temperature 37°C, where the offspring of *Microsporidia MB* negative mothers had three times higher pupation rate compared to the offspring of *Microsporidia MB* positive mothers [37°C; MB-mothers : 40.9 (0.29-0.53) %] (Tukey test at 37°C: p_MB-/MB+_ < 0.001; other Tukey tests: p > 0.05; *χ*^2^= 11.26, df = 1, p < 0.003).

**Figure 1:**
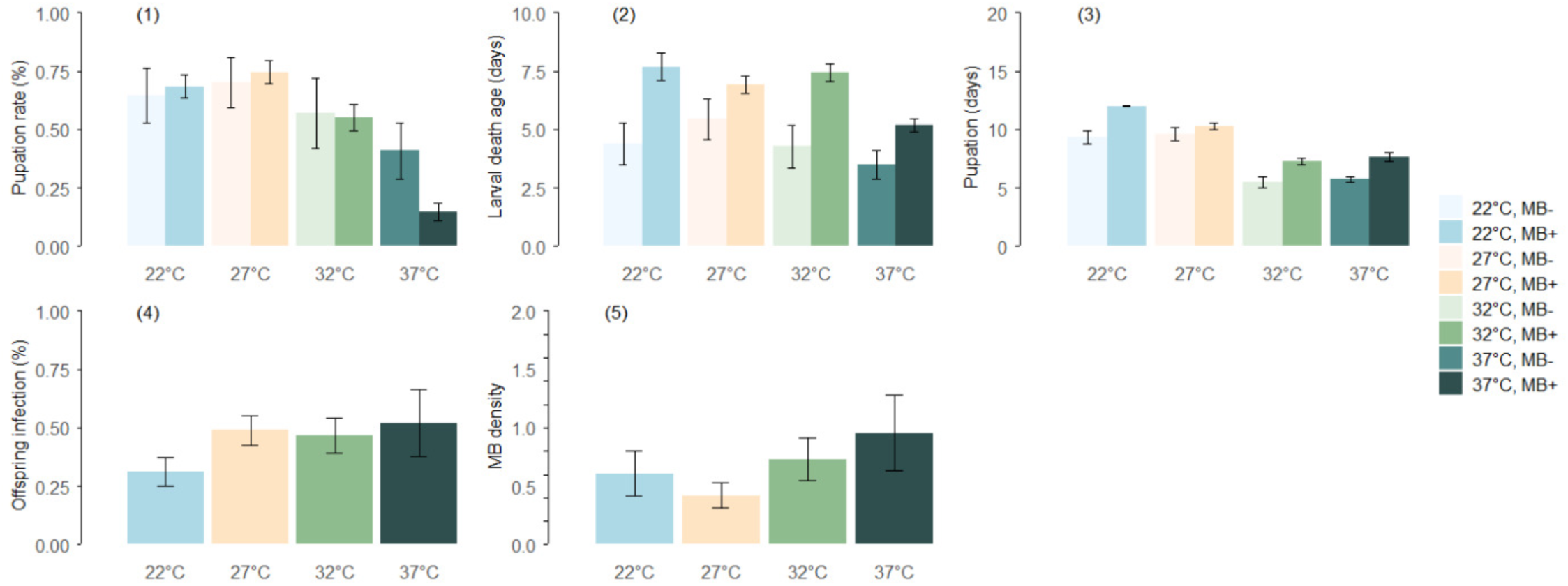
Panel representing the average (**1**) Pupation rate, (**2**) Age at death, (**3**) Larval development time, (**4**) Infection rate, (**5**) M*icrosporidia MB* intensity (log transformed ratio *Microsporidia MB* 18S/*Anopheles* S7 (ratio_MB18S/S7_) for better data visualization) in ofspring coming from non-infected (MB-, lighter colours) and infected mothers (MB+, darker colours) reared in 4 temperature treatments: 22°C (blue bars), 27°C (tan bars), 32°C (green bars) or 37°C (emerald bars). Average pupation rates are calculated out of the total count of ofspring, average larval death ages are calculated for larvae that died, average times to pupation were calculated for all individuals that pupated, ofspring infection rates were calculated for all individuals coming from infected mothers, and average *Microsporidia MB* intensities were calculated for all infected ofspring. The error bars show conûdence intervals.

#### Larval death age

larvae that did not pupate died on average after 6.1 (5.79-6.33) days post-hatching from their eggs. Larvae died around two days earlier when reared at temperature 37°C compared to all other temperatures [22°C: 7.1 (6.17-8.02) days; 27°C: 6.6 (5.93-7.35) days; 32°C: 7.1 (6.53-7.60) days; 37°C: 5.0 (4.69-5.27) days; **(Figure 1.2)**] (Tukey for dead larvae: p_27°C-37°C_, p_22°C-37°C_ and p_32°C-37°C_ < 0.001; *χ*^2^= 39.79, df = 3, p < 0.001). *Microsporidia MB* negative mothers had offspring dying two days earlier on average than the offspring from the *Microsporidia MB* positive mothers regardless of the temperature treatments [MB-mothers: 4.2 (3.62-4.86) days; MB+ mothers: 6.4 (6.07-6.66) days] (*χ*^2^= 18.66, df = 1, p < 0.001), except at 27°C where we found no significant differences between the mothers’ groups (Tukey 27°C: p_MB=/MB-_ > 0.05); *χ*^2^= 9.98, df = 3, p = 0.02).

#### Mean larval development time

The average larval development time (time between hatching and pupation) for the larvae that pupated was 9.6 (9.41-9.80) days. *Microsporidia MB* negative mothers had offspring pupating almost two days faster than offspring from *Microsporidia MB* positive mothers [MB-mothers: 8.1 (7.67-8.47) days; MB+ mothers: 9.9 (9.71-10.13) days] (*χ*^2^= 56.87, df = 1, p < 0.001). However, being *Microsporidia MB* positive due to maternal transmission increased the chance to pupate more than one day faster compared to Microsporidia MB negative larvae coming from infected mothers [MB+ offspring: 9.1 (8.85-9.40) days; MB-coming from MB+ mothers: 10.2 (9.93-10.52) days] (*χ*^2^= 50.92, df = 2, p < 0.001), thanks to a significant difference at 27°C [MB-offspring from MB+ mothers: 11.1 (10.66-11.46) days; MB+ offspring: 9.4 (9.06-9.83) days] (Tukey: p_MB+mother only-MB+offspring_ = 0.02; *χ*^2^= 21.92, df = 6, p = 0.001). At 22°C, 32°C, and 37°C, all larvae coming from *Microsporidia MB* infected mothers had similar development times (all Tukey test p > 0.05). The development time was negatively correlated with increasing temperature. At 32°C and 37°C, the time to pupation was 36 % lower compared to the 27°C treatment and almost 50 % lower compared to the 22°C treatment [22°C: 11.6 (11.29-11.88) days; 27°C: 10.1 (9.86-10.36) days; 32°C: 7.1 (6.89-7.42) days; 37°C: 6.96 (6.62-7.30) days] (Tukey for larvae that pupated: for all p < 0.001, except p_32°C-37°C_ > 0.05; *χ*^2^= 137.95, df = 3, p < 0.001). In offspring coming from *Microsporidia MB* negative mothers, the time to pupation did not vary between 22°C and 27°C (Tukey: p_22°C-27°C_ > 0.05), but we observed a two days difference in offspring coming from Microsporidia MB positive mothers reared in these temperatures [22°C: 12.0 (11.70-12.31) days; 27°C: 10.2 (9.93-10.48) days; 32°C: 7.4 (6.89-7.42) days; 37°C: 7.6 (7.25-8.04) days, **(Figure 1.3)**] (Tukey: p_22°C-27°C_ < 0.001; *χ*^2^= 21.89, df = 3, p < 0.001).

#### Infection rate in offspring

The infection rate in the 654 larvae coming from *Microsporidia MB* positive mothers was 43.0 (39.17-46.77) %. The infection rate was 32% lower at 22°C compared to the infection rate at 27°C [22°C: 30.8 (24.45-37.22) %; 27°C: 48.7 (42.43-5507) %; 32°C: 46.7 (39.06-54.28) %; 37°C: 52.1 (37.95-66.21)%] **(Figure 1.4)** (Tukey: estimate_27°C-22°C_ = 0.68, p_27°C-22°C_ = 0.001; p_27°C-32°C_ and p_27°C-37°C_ > 0.05; *χ*^2^= 9.99, df = 1, p = 0.001). The infection rate in the offspring was positively correlated with the intensity of *Microsporidia MB* in the mothers (*χ*^2^= 69.55, df = 1, p < 0.001).

#### *Microsporidia MB* intensity in offspring

the average relative *Microsporidia MB* intensity in *Microsporidia MB* positive offspring was 1.7 (1.19-2.23) ratio *Microsporidia MB* 18S/*Anopheles* S7 (ratio_MB18S/S7_). The temperature 27°C had the lowest *Microsporidia MB* intensity in offspring and significantly different from the temperature 37°C [22°C: 2.2 (0.34-4.01) ratio_MB18S/S7_; 27°C: 1.0 (0.55-1.43) ratio_MB18S/S7_; 32°C: 2.2 (1.27-3.11) ratio_MB18S/S7_; 37°C: 2.55 (1.28-3.75) ratio_MB18S/S7_; (Tukey: p_22°C-37°C_ < 0.001; p_27°C-37°C_ = 0.001; *χ*^2^= 19.40, df =3, p < 0.001)] (**Figure 1.5)**. Generally, offspring with high *Microsporidia MB* intensity pupated faster (*χ*^2^= 6.56, df = 1, p = 0.01), but this relationship was dependent on temperature (p_22°C-27°C_ < 0.001; p_27°C-37°C_ = 0.06; p_27°C-32°C_ = 0.001; *χ*^2^= 17.45, df = 3, p < 0.001); at 27°C, pupating 1 day faster increased the average *Microsporidia MB* intensity by 45 % (y(_27_) = 9.7 - 0.467x, r^2^ = 0.12); however at 22°C, a delay of 1 day in pupation time increased the *Microsporidia MB* intensity by 45% (y(_22_) = 11.4 + 0.311x, r^2^ = 0.13). At 27°C and 37°C, the development time was negatively correlated to *Microsporidia MB* intensity in the offspring (y(_37_) = 7.75 - 0.252x, r^2^ = 0.23). At 32°C, *Microsporidia MB* intensity in the offspring was not influenced by the development time (y(_32_) = 7.33 + 0.00623x, r^2^ < 0.01). Additionally, when mothers had a transmission rate higher than 50%, the *Microsporidia MB* intensity in offspring was higher [between 0 and 33% (low): 0.08 (0.021-0.144) ratio_MB18S/S7_; between 33% and 66% (medium): 0.33 (0.216-0.445) ratio_MB18S/S7_; between 66% and 99% (high): 0.85 (0.734-0.981) ratio_MB18S/S7_(Tukey: p_low-medium_ and p_low-high_ < 0.001, p_medium-high_ = 0.001; *χ*^2^= 162.66, df = 2, p < 0.001)].

### 2) Modelling the *Microsporidia MB* dissemination potential

We observed significant variations in the time required to reach a target population of 1000 *Microsporidia MB* positive mothers from a starting population of 10 mothers depending on temperature under Monte Carlo simulations and fecundity rate of 99 offspring per mother mosquitoes (**Figure 2.1**). As temperature increased, the mean age at which offspring pupate declined, while the probability of successful pupation did rise.

**Figure 2:**
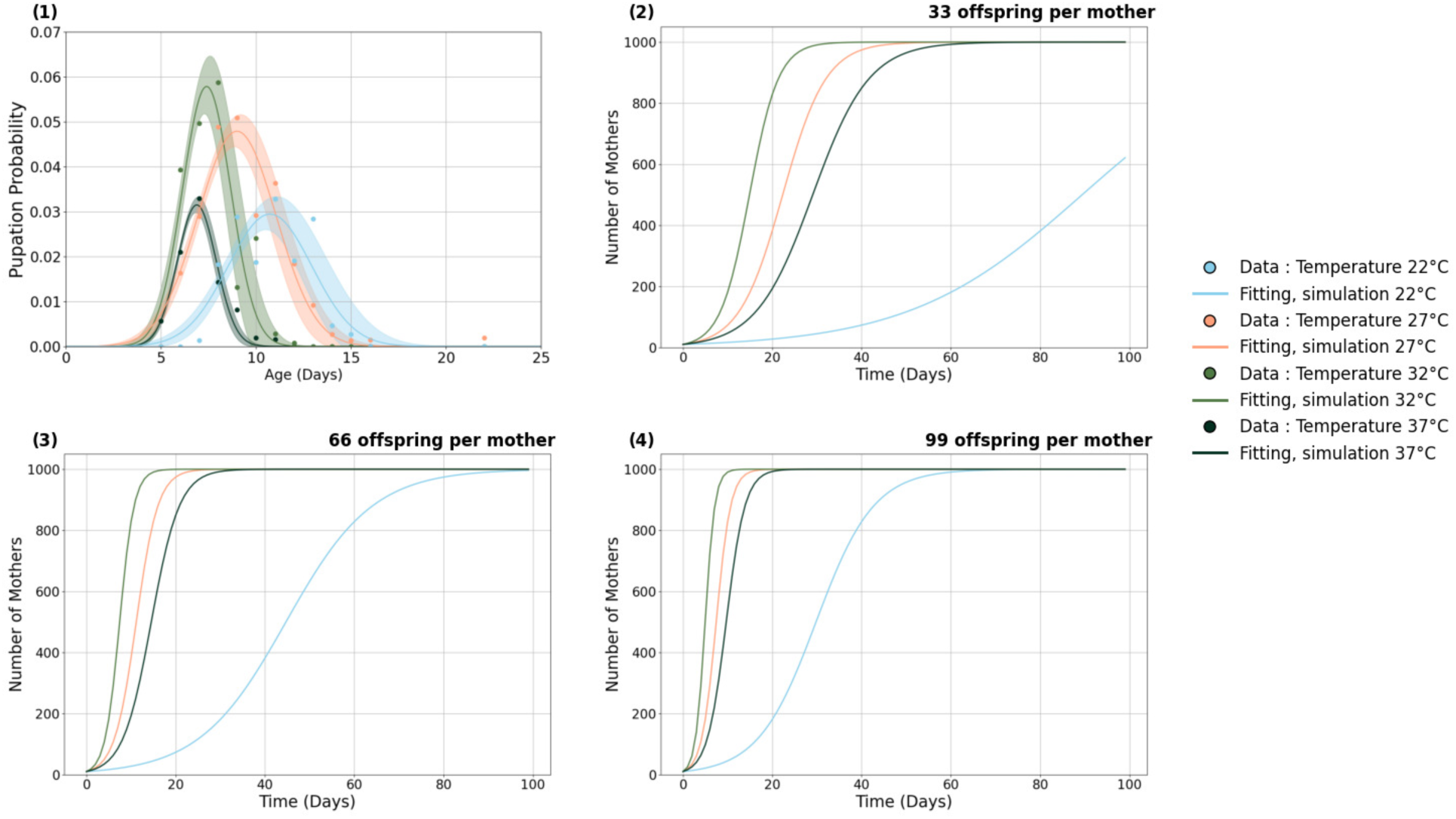
Panel representing the effects of 4 different temperature treatments: 22°C (blue color), 27°C (tan color), 32°C (green color) or 37°C (emerald color) on (**1**) The probabilities 9(T, x) modelled using a gaussian function that an offspring is infected, survives to age x, and pupates at age x and (**3**), (**4**), (**5**) represent the population growth of infected offspring starting with an initial population of 10 mothers and progressing over a 100-day period across the 4 four temperature treatments considering different fecundity levels; 33, 66 and 99 offspring per mother, respectively. Each growth curve incorporates a sex ratio of 0.5 (indicating an equal proportion of female offspring), a generation cycle and is capped by a carrying capacity K = 1000, illustrating how temperature affects the speed and likelihood of reaching the target population over time.

The probability that an offspring is infected, survives to age x, and pupates at age x given temperature T was modelled as a function of fitted parameters: A (scaling factor), mu (µ) (mean age of pupation), and sigma (σ) (spread of the pupation age). This probability combined the likelihood of infection at temperature T, the chance of survival to age x given infection, and the probability of pupating at that specific age. The model’s accuracy was validated by high R^2^ values, especially at higher temperatures, indicating a strong fit to the observed data. At 22°C, the normalization constant A was 0.02942, with a mean pupation age mu of 10.73 and a standard deviation sigma of 2.20, resulting in a mean squared error (MSE) of 0.00003 and a R^2^ value of 0.794, indicating a moderate fit. At 27°C, A increased to 0.04785, mu decreased to 8.99, and sigma was 2.06, with an MSE of 0.00003 and a stronger fit at R^2^ = 0.917. At 32°C, A further increased to 0.05782, mu dropped to 7.40, and sigma narrowed to 1.27, producing an MSE of 0.00005 and an R^2^ of 0.874. Finally, at 37°C, A was 0.03147, mu was 6.88, sigma was 1.02, with the best fit demonstrated by an MSE of 0.00000 and a R^2^ of 0.970.

The population growth for infected offspring across different temperatures and fecundity levels revealed that both factors significantly impact growth rates and time to reach population of 1000 *Microsporidia MB* positive mothers. The optimum temperatures 27°C and 32°C consistently led to rapid population growth, with 1000 *Microsporidia MB* positive mothers reached within 15–48 days across all fecundity levels. Specifically, at a fecundity of 33 offspring per mother **(Figure 2.2)**, populations at 27°C and 32°C reach 1000 *Microsporidia MB* positive mothers by approximately day 48 and day 35, respectively, while at 37°C, it was estimated to take around 62 days. In contrast, at 22°C, the population did not reach 1000 mothers *Microsporidia MB* positive within the 100-day period. Increasing fecundity to 66 offspring per mother did accelerate growth further, with populations at 27°C, 32°C, and 37°C reaching 1000 *Microsporidia MB* positive mothers by approximately day 25, 20, and 35, respectively **(Figure 2.3).** At 22°C, the average time to reach the target population was 100 days. At the highest fecundity level of 99 offspring per mother **(Figure 2.4)**, the population growth was extremely rapid, with 27°C and 32°C reaching 1000 *Microsporidia MB* positive mothers by day 17 and 15, respectively, and 37°C by day 25. At 22°C, the population reached 1000 *Microsporidia MB* positive mothers by approximatively 65 days.

To further clarify the effects of offspring numbers per mother, we conducted a Monte Carlo simulation was conducted with 1000 iterations, accounting for 10% variability in offspring number (Supplementary information, Figure A). This analysis provides the average establishment time across 1000 simulations for each temperature. Results indicate that temperatures of 27°C and 32°C remain optimal for the establishment of *Microsporidia MB*, regardless of fluctuations in the offspring count per mother.

## Discussion

The optimum temperature for maintaining the symbiont *Microsporidia MB* intensity and supporting *Microsporidia MB* infected *Anopheles* mosquito population growth was estimated to be 32°C. This contrasts with the optimum temperature for rearing *Anopheles* mosquitoes often considered to be 27°C. Despite a recorded higher survival rate for *Microsporidia MB* infected (and un-infected mosquitoes) at 27°C, the rate of population growth for a *Microsporidia MB* infected *Anopheles* population is optimal at 32°C as a result of a shorter larval development time and higher *Microsporidia MB* intensity in the offspring. Overall, temperatures ranging from 27°C to 32°C and closer to 32°C can be considered most efficient rearing temperatures for symbiont-based malaria control tool development by mass rearing *Microsporidia MB* infected *Anopheles* before field releases (note that we did not yet evaluate the malaria blocking potential at different temperatures). Our experiments were conducted on wild-caught *Anopheles* mosquitoes and their offspring which have relatively low fecundity (an estimated average of 33-66 offspring per female) relative to lab-adapted and reared colonies of *Anopheles* mosquitoes (24). Our findings underscore the critical interplay between air temperature and mosquito fecundity in determining the population growth of *Microsporidia MB* positive mothers. The results suggest that optimizing these factors could enhance the efficiency of *Microsporidia MB* as a potential vector control strategy. Additionally, the strong correlation coefficients (R² values ranging from 0.79396 to 0.97048) further support the reliability of our simulations, indicating that the logistic growth model effectively captured the dynamics of *Microsporidia MB* positive population changes under varying environmental conditions.

Mosquitoes are poikilotherms and their life history characteristics highly depend on the surrounding temperatures. These characteristics include longevity, fecundity, biting rate and development of immature stages of mosquitoes (25). Our study confirms previous studies about warmer temperatures reducing the pupation rate and reducing the larval development time, regardless of the infection to *Microsporidia MB* (43). In addition, we found that larvae coming from *Microsporidia MB* infected mothers took almost two days longer time to develop compared to larvae coming from *Microsporidia MB* negative mothers. However, when the offsprings of *Microsporidia MB* positive mothers were separated between infected and non-infected larvae in the statistical model, we realized that *Microsporidia MB* positive offspring were overall developing 1 day faster than *Microsporidia MB* negative offspring coming from *Microsporidia MB* infected mothers thanks to a significant difference in development at 27°C. This result is similar to earlier reports performed in the laboratory and semi-field conditions where temperature was not controlled (7,26,27).

The occurrence of some genera of microsporidians (*Enterocytospora, Microsporidium* and *Vairimorpha*) are positively correlated with increase in environmental temperature (8). Similarly, we observed that the prevalence of *Microsporidia MB* was increased at higher temperatures. A higher intensity of *Microsporidia MB* infection was recorded at higher temperatures; therefore, our results indicate that transmission rate which is *Microsporidia MB* intensity dependent is influenced by temperature. Likewise in *Drosophila*, higher temperatures result in greater proliferation of the bacteria *Wolbachia* (28,29) and higher cost of the pathogenic variants such as *wMelOctoless* and *wMelPop* in *D. melanongaster* (28). Lower development temperatures lead to lower titres of *wYak* in *Drosophila yakuba*, reducing the chances of vertical transmission. In our study, the coolest rearing temperature was 22°C and resulted in the “loss” of *Microsporidia MB* infection in offspring after vertical transmission from a mother to her offspring. The teneral reserve might influence the loss of *Microsporidia MB*; when the temperatures are cooler, the development time is slower and mortality timing in larvae is late during the development, the chance of survival for *Microsporidia MB* are lower and the symbiont might use the horizontal route of transmission to ensure its survival (30–32). This shows the importance of elucidating the mechanisms of that symbiosis and better understand the benefits brought by the symbionts and what it would mean for *Anopheles* mosquitoes to losing them. The effect of temperature on the strategy used by the symbiont *Microsporidia MB* to disseminate vertically or horizontally was not studied here, but we hypothesize that extreme cool temperatures which ensure a slow larval death and a long development until pupation might potentially be favourable for the formation of *Microsporidia MB* spores (33,34). In our previous studies we have not observed larvae-to-larvae transmission, however we cannot rule out that combinations of physiological stressors and less-than-optimal rearing conditions could promote horizontal transmission at the larval stage. In other microsporidian systems, such as *Edhazardia aedis* the impact of resource availability on the transmission route showed that there is an ecological trade-off between transmission modes resulting in higher horizontal transmission potential when mosquito larvae are reared under low-food regime, a stress that increases the larval development time (35,36).

We observed the highest *Microsporidia MB* infection intensity and prevalence at 37°C, which may suggest that higher temperatures result in a higher growth rate of *Microsporidia MB* in *Anopheles*. A higher *Microsporidia MB* growth rate would result in a higher infection intensity but may also enable *Microsporidia MB* to decrease the probability that hosts either lose or clear their *Microsporidia MB* infection (resulting in higher observed *Microsporidia MB* prevalence). However, in our study the highest intensity of infection and mortality of *Microsporidia MB* infected larvae was observed at 37°C, an extremely hot temperature for mosquito rearing. 37°C is considered an upper thermal limit for adult mosquito development (37–39), however, heat tolerance at the larval stage is higher compared to the adult stage. It is unclear why we observed such a low pupation rate for offspring coming from *Microsporidia MB* infected mothers compared to those coming from *Microsporidia MB* uninfected mothers. One hypothesis is that 37°C potentially is a thermal limit for *Microsporidia MB*; the intensity in *Microsporidia MB* infected offspring increases but the heat kills some of the symbiont population, hence it becomes toxic or pathogenic to the host and result in a high mortality rate in *Microsporidia MB* infected mosquitoes (8). The presence of the symbiont might reduce the heat tolerance of mosquitoes and disrupt the host’s ability to cope with thermal stress, as observed in *Daphnia magna* with the obligate bacterial pathogen *Pasteuria ramosa* (40). In fact, the thermal mismatch hypothesis predicts that cooler-adapted hosts faced with warmer-adapted parasites are more at risk to succumbing to infection in warmer temperature (41). This hypothesis warrants for future studies on the sensitivity of *Microsporidia MB* to temperature.

It is also conceivable that higher temperatures, which dramatically increase larval death, favour the formation of spores and horizontal transmission. Assuming that infection via spores results in some mortality, this could be an alternative explanation for the increase in *Microsporidia MB* prevalence and *Anopheles* mortality in hot larval rearing conditions. The effect of heat stress on different types of *Wolbachia* was observed by Ross *et al.,* (42), and an increase in rearing temperature from 26° C to 37°C for the rearing of all life stages of *Wolbachia*-infected *Aedes aegypti* mosquitoes reversed the infection and eventual transmission of the endosymbiont to the next generation of mosquitoes.

It is well known that environmental conditions experienced during larval development influence adult mosquito life-history traits. Variation in larval habitats quality carry over to affect adult life history (19,43). In this study we did not focus on the adult stage of mosquitoes, hence it limits the predictions about population growth potential of mosquitoes reared in different temperatures. The survival rate is well-underestimated in our study due to a data collection focused on juvenile development only, even though higher temperatures have been reported to adversely affect the longevity of blood-fed mosquitoes and their fecundity (37). High temperature also reduces the body size and decrease the hatch rate (5,17,18,44). Lower body size live longer in some environments but not in others, depending on the food resources and temperature during larval development (19). These two rearing parameters interact also in complex ways to influence vector competence for malaria, with opposite relationship between temperature and competence depending on larval food intake (5,45). We hypothesize that similar complex relationship exists for *Microsporidia MB* competence and warrant further investigation. Also, even though the intensity of *Microsporidia MB* is influenced by nutrition (23,26), this study assumed similar larval competition for food in all temperature treatments. As density-dependent competition and temperature are known to interact to influence mosquito survival, which trade-off with offspring number (35,36), there is a need to better characterize the impact of nutrition intake in *Microsporidia MB* infected mosquitoes. This is even more important as we report the influence of temperature on *Microsporidia MB* spread depending on the number of offspring generated: with 33 offspring per mother, 22°C prevent the growth of *Microsporidia MB* infected populations. Only when the offspring count is high (99 offspring per mother) do temperatures of 22°C and 37°C favour the rapid establishment of *Microsporidia MB*.

## Conclusion

The optimal rearing temperature for *Anopheles* mosquitoes is known to be 27°C; however, that reference is not the optimal rearing temperature for the dissemination of the symbiont *Microsporidia MB*. Despite the similar offspring infection rate between 27°C and 32°C, plus a better mosquito pupation rate at 27°C compared to 32°C, the symbiont intensity in *Microsporidia MB* positive offspring was higher at 32°C and the development time at the larval stage quicker at 32°C compared to 27°C. This work confirms the previous field data collection and modelling prediction highlighting the preference of the symbiont for warmer temperatures, despite a potentially high mosquito mortality rate. It also helps further identify regions where climate is favorable to the spread of *Microsporidia MB* infected mosquito populations for better malaria control. In this paper, we highlight the importance of considering environmental factors in the efficacy assessment of microbes-based malaria control tools to help disseminate *Microsporidia MB* positive mosquitoes in nature and better understand the natural prevalence and spread of the symbiont in different climatic regions.

## Material and methods

### 1) Sample collection, rearing process and quantification of *Microsporidia MB*

1539 larvae used in this study were obtained from 17 wild caught gravid *Anopheles arabiensis* female collected in Kigoche village (00°34ʹS, 34°65ʹ E) via mouth aspiration in the Ahero irrigation scheme, Kenya and transported to the icipe-Duduville campus in Nairobi, Kenya. These females laid on average 65.8 (standard deviation: 26.36) eggs and were selected over three different collection timepoints : September 2022 (5 females, offspring n = 404), November 2022 (5 females, offspring n = 511) and July 2023 (7 females, offspring n = 624), placed in 1.5ml micro-centrifuge tubes containing 1cm by 1cm Whatman filter paper to allow egg laying following the methods described in (7,23) and, after oviposition, screened for species ID (46) and the presence of *Microsporidia MB* (7) using PCR.

Eggs from *Microsporidia MB* positive and negative mothers were separated in larval trays with around 300 ml of deionised water to hatch. F1 offspring were reared on our controlled Tetramin baby fish food regime from hatching until pupation and L1 larvae were randomly split in four temperature treatments: 22°C ± 1 (n= 398), 27°C ± 1 (n= 393), 32°C ± 1 (n= 351), or 37°C ± 1 (n= 397). We also used 3 female lines from our susceptible *An. arabiensis* insectary colony reared in the same condition as control. We measured larvae daily mortality and individual pupation timing and rate.

All pupae were screened for *Microsporidia MB* quantification using PCR (6,7). We measured the *Microsporidia MB* infection rate in the mothers and offspring as well as *Microsporidia MB* density. The ammonium acetate protein precipitation method was used for DNA extraction. Partial *Microsporidia MB* 18s gene region from each DNA sample was amplified using specific 18s primers (MB18SF: CGCCGG CCGTGAAAAATTTA and MB18SR: CCTTGGACGTG GGAGCTATC) (6,7,23). The gene was then amplified in an 11µl reaction volume of a mixture containing 0.5µl of 5pmol/µl reverse and forward primers, 2µl HOTFirepol Blend Master Mix Ready-To-Load (Solis Biodyne, Estonia), 6µl of nuclease-free PCR water and 2µl of DNA template. The amplification was achieved under the following conditions: initial denaturation at 95 °C for 15 min, denaturation at 95 °C for 1 minute for 35 cycles, annealing at 62 °C for 30 s, a further extension for 30 s at 72 °C, and finally, final elongation for 5 min at 72 °C. To quantify the level of infection, samples positive for *Microsporidia MB* were subjected to relative qPCR analysis using MB18SF/MB18SR primers normalised with the reference host-keeping gene for the *Anopheles* ribosomal s7 gene (S7F: TCCTGGAGCTGGAGATGAAC and S7R: GACGGGTCTGTACCTTCTGG). The qPCR reaction mixture consisted of 11µl reaction volume containing 0.5µl of 5pmol/µl reverse and forward primers, 2µl HOT FIREPol® EvaGreen® 416 HRM no ROX Mix Solis qPCR Master mix (Solis Biodyne, Estonia), 6µl of nuclease-free PCR water and 2µl of DNA template. The amplification was achieved under the following conditions: initial denaturation at 95 °C for 15 min, denaturation at 95 °C for 1 minute for 35 cycles, annealing at 62 °C for 60 s, and a further extension for 45 s at 72 °C. The PCR was carried out in a proflex cycler, and the qPCR was carried out in a MIC qPCR cycler (BioMolecular Systems, Australia). The MB18SF/MB18SR primers were used to confirm samples with the characteristic *Microsporidia MB* melt curve (7).

We analysed the pupation rate and age at death using Mixed-Effects Cox Models and the R “coxme” package (47). The mean development time for the pupated larvae was analysed using the linear mixed-effects model using the “lme4” package. We analysed the infection rate and *Microsporidia MB* intensity using binomial and gaussian logistic mixed-effect model (GLMMs) and glmmTMB package. In all models, the temperature treatments, the mothers’ infection status, and their interactions were included as fixed terms, and the time of capture in the field was included as a random effect. In addition, the development time model also looked at the interaction between temperature treatments and infection status in offspring (*Microsporidia MB* negative offspring coming from un-infected colonized mothers, *Microsporidia MB* positive offspring coming from field-collected infected mothers and *Microsporidia MB* negative coming from field collected infected mothers). Individuals that pupated were excluded from the age-at-death analysis. Individuals who died were excluded from the development time and infection status analysis. The *Microsporidia MB* intensity analysis (log transformed for better data visualisation) excluded uninfected pupae, and we used temperature treatments and transmission groups (0-33%, 33-66%, or 66-99% transmission from mothers to offspring) in interaction in the model. We used the Tukey post-hoc test and “means” function to perform multiple comparisons among the infection status and temperature treatments (48). Statistical analysis was performed using R statistical software version 4.1.2 and R Studio (49).

### 2) Modelling the *Microsporidia MB* dissemination potential

To express the probability that an L1 offspring coming from MB+ mother is infected, survives to age x, and pupates at age x given temperature T, we combined the conditional probabilities:

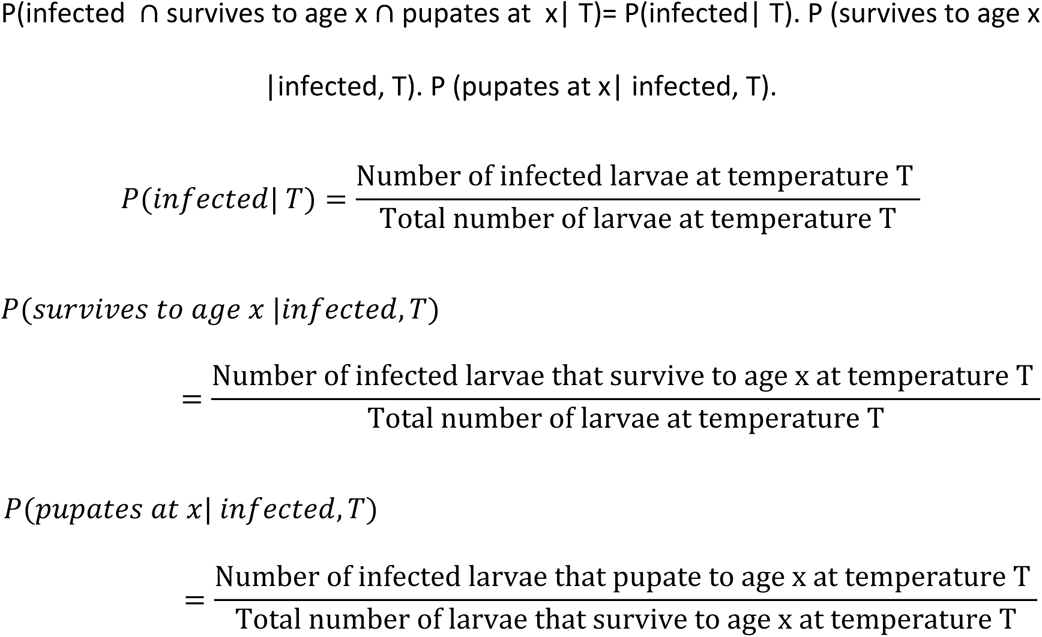

Using then Gaussian function, the probability is given by:

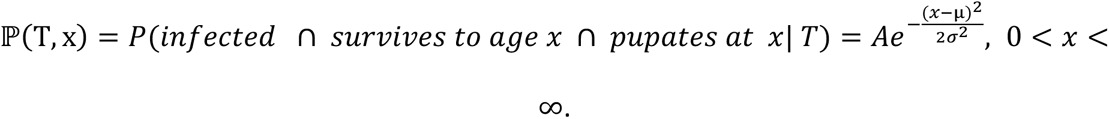

This formula considers the conditional dependencies based on infection status and temperature, providing a logical path to estimate the combined probability.

In the following section of the methodology, we employed the logistic growth model to simulate the dissemination potential of *Microsporidia MB* in mosquito populations, specifically tracking the growth of infected mothers under different temperature conditions. The logistic growth equation,

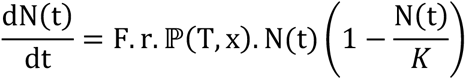

models the population growth of infected individuals, where N(t) is the number of MB+ individuals at time t, F represents the fecundity, r the sex ratio, ℙ(T, x) the probability of infection, survival, and pupation under temperature T, and K the carrying capacity. The solution to this equation,

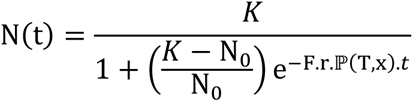

enabled us to estimate the rate at which the population of MB+ offspring increases from an initial population of 10 MB+ mothers, with the goal of reaching a target population of 1000 *Microsporidia MB* positive individuals.

In our deterministic simulation, parameters such as: F (fecundity), r (sex ratio), K (carrying capacity), and N_"_(initial population) remained constant. Fecundity was set at three fixed rates (33, 66, or 99 viable eggs per mother) based on observed averages, providing a baseline for population growth under stable conditions. Details of the stochastic simulation are provided in the supplementary material.

To implement this methodology, we used Python for all data processing, simulations, and statistical computations. Python’s libraries, including *numpy* for numerical operations, *scipy* for probability computations and fitting, and *matplotlib* for visualization, were integral to generating plots, calculating probabilities, and fitting model parameters.

## Availability of data and materials

All datasets generated, used and analyzed in this study are included in this published article.

## Acknowledgements

We acknowledge the Insectary team led by Milca Gitau, Jeniffer Thiong’o and Peris Wambui for provision of larvae rearing materials and assistance in the rearing process. The field team (Robison Kisero and Gerald Ronoh) for assistance during field collection of gravid mothers. The project administrator Faith Kyengo for facilitating all the project activities.

## Author Contributions

Fidel Gabriel Otieno, Conceptualization, Data curation, Methodology and Investigation, Validation and Visualisation of results, Writing of the original draft, review and editing ∣ Priscille Barreaux, Conceptualization, Supervision, Formal analysis, Validation and Visualization of results, Writing of the original draft, review and editing ∣ Affognon Steeven Belvinos, Formal analysis, Validation and Visualization of results, Writing of the original draft, review and editing ∣ Edward Edmond Makhulu, Data curation, Methodology, Investigation ∣ Thomas Ogao Onchuru, Methodology, Investigation, Supervision, review and editing∣ Anne Wambui Wairimu, Investigation, Data curation, Methodology ∣ Stancy Mandere Omboye, Investigation, Data curation, Methodology ∣ Cynthia Nyambura King’ori, Investigation, Data collection, Methodology ∣ Sokame Bonoukpoè Mawuko, Formal analysis, Validation and Visualization of results, Writing of the original draft, review and editing ∣ Anthony Nyamache Kebira, Conceptualization, Supervision ∣ Jeremy Keith Herren, Conceptualization, Methodology and Investigation, Supervision, Funding acquisition, Resources, Writing of the original draft, review and editing.

## References

1. Filho, W. L., May, J., May, M., & Nagy, G. J. (2023). Climate change and malaria: some recent trends of malaria incidence rates and average annual temperature in selected sub-Saharan African countries from 2000 to 2018. Malaria Journal, 22(1). 10.1186/s12936-023-04682-4

2. World Health Organization. (2022, December 8). World malaria report 2022. www.who.int. https://www.who.int/publications/i/item/9789240064898

3. Ma, C.-S., Zhang, W., Peng, Y., Zhao, F., Chang, X.-Q., Xing, K., Zhu, L., Ma, G., Yang, H.-P., & Rudolf, V. H. W. (2021). Climate warming promotes pesticide resistance through expanding overwintering range of a global pest. Nature Communications, 12(1). 10.1038/s41467-021-25505-7

4. World Health Organization. (2023). WHO malaria policy advisory group (MPAG) meeting report, 18–20 April 2023. World Health Organization.

5. Barreaux, A. M. G., Priscille Barreaux, Thievent, K., & Koella, J. C. (2016). Larval environment influences vector competence of the malaria mosquito. PubMed, 7, 8–8. 10.5281/zenodo.10798340

6. Nattoh, G., Maina, T., Makhulu, E. E., Mbaisi, L., Mararo, E., Otieno, F. G., Bukhari, T., Onchuru, T. O., Teal, E., Paredes, J., Bargul, J. L., Mburu, D. M., Onyango, E. A., Magoma, G., Sinkins, S. P., & Herren, J. K. (2021). Horizontal transmission of the symbiont *Microsporidia MB* in *Anopheles arabiensis*. Frontiers in microbiology, 12. 10.3389/fmicb.2021.647183

7. Herren, J. K., Mbaisi, L., Mararo, E., Makhulu, E. E., Mobegi, V. A., Butungi, H., Mancini, M. V., Oundo, J. W., Teal, E. T., Pinaud, S., Lawniczak, M. K. N., Jabara, J., Nattoh, G., & Sinkins, S. P. (2020). A microsporidian impairs *Plasmodium falciparum* transmission in *Anopheles arabiensis* mosquitoes. Nature communications, 11(1). 10.1038/s41467-020-16121-y

8. Artur Trzebny, Olena Nahimova, & Miroslawa Dabert. (2024). High temperatures and low humidity promote the occurrence of microsporidians (*Microsporidia*) in mosquitoes (*Culicidae*). Parasites & Vectors, 17(1). 10.1186/s13071-024-06254-0

9. Willis, A. R., & Reinke, A. W. (2022). Factors that determine *Microsporidia* infection and host specificity. Experientia supplementum, 91–114. 10.1007/978-3-030-93306-7_4

10. Chakrabarti, S. B. (2016). Influence of temperature and relative humidity in infection of *Nosema bombycis* (Microsporidia: *Nosematidae*) and cross-infection of *N. mylitta* on growth and development of *Mulberry silkworm*, Bombyx mori. International Journal of Industrial Entomology and Biomaterials, 17(2), 173–180. https://koreascience.kr/article/JAKO200811237154541.page

11. Cali, A., & Takvorian, P. M. (2014). Developmental morphology and life cycles of the Microsporidia. 71–133. 10.1002/9781118395264.ch2

12. Chandrasegaran, K., Lahondère, C., Escobar, L. E., & Vinauger, C. (2020). Linking mosquito ecology, traits, behavior, and disease transmission. Trends in parasitology, 36(4), 393–403. 10.1016/j.pt.2020.02.001

13. Reinhold, J., Lazzari, C., & Lahondère, C. (2018). Effects of the environmental temperature on *Aedes aegypti* and *Aedes albopictus* mosquitoes: A review. Insects, 9(4), 158. 10.3390/insects9040158

14. Paaijmans, K. P., & Thomas, M. B. (2011). The influence of mosquito resting behaviour and associated microclimate for malaria risk. Malaria journal, 10(1). 10.1186/1475-2875-10-183

15. Martin, L. E., & Hillyer, J. F. (2024). Higher temperature accelerates the aging-dependent weakening of the melanization immune response in mosquitoes. PLOS Pathogens,20(1),e1011935–e1011935. 10.1371/journal.ppat.1011935

16. Maria Cecilia Mancini, Ant, T. H., Herd, C. S., Gingell, D. D., Murdochy, S. M., Enock Mararo, & Sinkins, S. P. (2020). High temperature cycles result in maternal transmission and dengue infection differences between *Wolbachia* strains in *Aedes aegypti*. 10.1101/2020.11.25.397604

17. Moller-Jacobs, L. L., Murdock, C. C., & Thomas, M. B. (2014a). Capacity of mosquitoes to transmit malaria depends on larval environment. Parasites & Vectors, 7(1). 10.1186/s13071-014-0593-4

18. Moller-Jacobs, L. L., Murdock, C. C., & Thomas, M. B. (2014b). Capacity of mosquitoes to transmit malaria depends on larval environment. Parasites & Vectors, 7(1). 10.1186/s13071-014-0593-4

19. Barreaux, A. M. G., Stone, C. M., Barreaux, P., & Koella, J. C. (2018). The relationship between size and longevity of the malaria vector *Anopheles gambiae (s.s.)* depends on the larval environment. Parasites & Vectors, 11(1). 10.1186/s13071-018-3058-3

20. Paaijmans, K. P., Blanford, S., Chan, B. H. K., & Thomas, M. B. (2011). Warmer temperatures reduce the vectorial capacity of malaria mosquitoes. Biology Letters, 8(3), 465–468. 10.1098/rsbl.2011.1075

21. Gimonneau, G., Bouyer, J., Morand, S., Besansky, N. J., Diabate, A., & Simard, F. (2010). A behavioral mechanism underlying ecological divergence in the malaria mosquito *Anopheles gambiae*. Behavioral ecology, 21(5), 1087–1092. 10.1093/beheco/arq114

22. Kikuchi, Y., Tada, A., Musolin, D. L., Hari, N., Hosokawa, T., Fujisaki, K., & Fukatsu, T. (2016). Collapse of insect gut symbiosis under simulated climate change. mBio, 7(5). 10.1128/mbio.01578-16

23. Makhulu, E. E., Onchuru, T. O., Gichuhi, J., Otieno, F. G., Wairimu, A. W., Muthoni, J. N., Koekemoer, L., & Herren, J. K. (2023). Localization and tissue tropism of the symbiont *Microsporidia MB* in the germ line and somatic tissues of *Anopheles arabiensis*. mBio. 10.1128/mbio.02192-23

24. Huang, J., Walker, E. D., Otienoburu, P. E., Amimo, F., Vulule, J., & Miller, J. R. (2006). Laboratory tests of oviposition by the african malaria mosquito, *Anopheles gambiae*, on dark soil as influenced by presence or absence of vegetation. Malaria journal, 5(1). 10.1186/1475-2875-5-88

25. Ciota, A. T., Matacchiero, A. C., Kilpatrick, A. M., & Kramer, L. D. (2014). The Effect of temperature on life history traits of *Culex* mosquitoes. Journal of medical entomology, 51(1), 55–62. 10.1603/me13003

26. Boanyah, G. Y., Koekemoer, L. L., Herren, J. K., & Bukhari, T. (2024). Effect of *Microsporidia MB* infection on the development and fitness of *Anopheles arabiensis* under different diet regimes. Parasites & Vectors, 17(1). 10.1186/s13071-024-06365-8

27. Godfrey Nattoh, Onyango, B., Edward Edmond Makhulu, Omoke, D., Lilian Mbaisi Ang’ang’o, Kamau, L., Maxwell Machani Gesuge, Ochomo, E., & Jeremy Keith Herren. (2023). *Microsporidia MB* in the primary malaria vector *Anopheles gambiae sensu stricto* is avirulent and undergoes maternal and horizontal transmission. Parasites & Vectors, 16(1). 10.1186/s13071-023-05933-8

28. Strunov, A. A., Ilinskii, Yu. Yu., Zakharov, I. K., & Kiseleva, E. V. (2013). Effect of high temperature on survival of *Drosophila melanogaster* infected with pathogenic strain of *Wolbachia* bacteria. Russian journal of genetics: applied research, 3(6), 435–443. 10.1134/s2079059713060099

29. Reynolds, K. T., Thomson, L. J., & Hoffmann, A. A. (2003). The effects of host age, host nuclear background and temperature on phenotypic effects of the virulent *Wolbachia* strain *popcorn* in *Drosophila melanogaster*. Genetics, 164(3), 1027–1034. 10.1093/genetics/164.3.1027

30. Dionysopoulou, N. K., Papanastasiou, S. A., Kyritsis, G. A., & Papadopoulos, N. T. (2020). Effect of host fruit, temperature and Wolbachia infection on survival and development of ceratitis capitata immature stages. PLOS ONE, 15(3), e0229727–e0229727. 10.1371/journal.pone.0229727

31. Ulrich, J. N., Beier, J. C., Devine, G. J., & Hugo, L. E. (2016). Heat Sensitivity of *wMel Wolbachia* during *Aedes aegypti* Development. PLOS Neglected Tropical Diseases, 10(7), e0004873. 10.1371/journal.pntd.0004873

32. Wiwatanaratanabutr, I., & Kittayapong, P. (2009). Effects of crowding and temperature on Wolbachia infection density among life cycle stages of *Aedes albopictus*. Journal of invertebrate pathology, 102(3), 220–224. 10.1016/j.jip.2009.08.009

33. Chrostek, E., Martins, N., Marialva, M. S., & Teixeira, L. (2021). *Wolbachia* -conferred antiviral protection is determined by developmental temperature. mBio, 12(5). 10.1128/mbio.02923-20

34. Lau, M.-J., Ross, P. A., Endersby-Harshman, N. M., & Hoffmann, A. A. (2020). Impacts of low temperatures on Wolbachia (Rickettsiales: *Rickettsiaceae*)-Infected *Aedes aegypti* (Diptera: *Culicidae*). Journal of medical entomology, 57(5), 1567–1574. 10.1093/jme/tjaa074

35. Zilio, G., Kaltz, O., & Koella, J. C. (2022). Resource availability for the mosquito *Aedes aegypti* affects the transmission mode evolution of a microsporidian parasite. Evolutionary ecology. 10.1007/s10682-022-10184-7

36. Zilio, G., Thiévent, K., & Koella, J. C. (2018). Host genotype and environment affect the trade-off between horizontal and vertical transmission of the parasite *Edhazardia aedis*. BMC Evolutionary Biology, 18(1). 10.1186/s12862-018-1184-3

37. Agyekum, T. P., Botwe, P. K., Arko-Mensah, J., Issah, I., Acquah, A. A., Hogarh, J. N., Dwomoh, D., Robins, T. G., & Fobil, J. N. (2021). A Systematic review of the effects of temperature on *Anopheles* mosquito development and survival: Implications for malaria control in a future warmer climate. International journal of environmental research and public health, 18(14), 7255. 10.3390/ijerph18147255

38. Carrington, L. B., Armijos, M. V., Lambrechts, L., Barker, C. M., & Scott, T. W. (2013). Effects of fluctuating daily temperatures at critical thermal extremes on *Aedes aegypti* life-history traits. PLoS ONE, 8(3), e58824. 10.1371/journal.pone.0058824

39. Lyons, C. L., Coetzee, M., Terblanche, J. S., & Chown, S. L. (2012). Thermal limits of wild and laboratory strains of two African malaria vector species, *Anopheles arabiensis* and *Anopheles funestus*. Malaria journal, 11(1). 10.1186/1475-2875-11-226

40. Hector, T. E., Sgrò, C. M., & Hall, M. D. (2019). Pathogen exposure disrupts an organism’s ability to cope with thermal stress. Global change biology, 25(11), 3893– 3905. 10.1111/gcb.14713

41. Hector, T. E., Hoang, K. L., Li, J., & King, K. C. (2022). Symbiosis and host responses to heating. Trends in ecology & evolution, 37(7), 611–624. 10.1016/j.tree.2022.03.011

42. Ross, P. A., Wiwatanaratanabutr, I., Axford, J. K., White, V. L., Endersby-Harshman, N. M., & Hoffmann, A. A. (2017). Wolbachia infections in *Aedes aegypti* differ markedly in their response to cyclical heat stress. PLOS Pathogens, 13(1), e1006006. 10.1371/journal.ppat.1006006

43. Christiansen-Jucht, C. D., Parham, P. E., Saddler, A., Koella, J. C., & Basáñez, M.-G. (2015). Larval and adult environmental temperatures influence the adult reproductive traits of *Anopheles gambiae s.s*. Parasites & Vectors, 8(1). 10.1186/s13071-015-1053-5

44. Tuno, N., Farjana, T., Uchida, Y., Iyori, M., & Yoshida, S. (2023). Effects of temperature and nutrition during the larval period on life history traits in an invasive malaria vector *Anopheles stephensi*. Insects, 14(6), 543. 10.3390/insects14060543

45. Araújo, M. da-Silva, Gil, L. H. S., & e-Silva, A. de-Almeida. (2012). Larval food quantity affects development time, survival and adult biological traits that influence the vectorial capacity of *Anopheles darlingi* under laboratory conditions. Malaria journal, 11(1). 10.1186/1475-2875-11-261

46. Santolamazza, F., Mancini, E., Simard, F., Qi, Y., Tu, Z., & della Torre, A. (2008). Insertion polymorphisms of SINE200 retrotransposons within speciation islands of *Anopheles gambiae* molecular forms. Malaria journal, 7(1). 10.1186/1475-2875-7-163

47. Jan, Erhardt, E. B., Dodd, A. B., Nathaniel, U., Wick, T. V., McQuaid, J. R., Ryman, S. G., Vakhtin, A. A., Meier, T. B., & Mayer, A. R. (2024). A cautionary tale on the effects of different covariance structures in linear mixed effects modelling of fMRI data. Human Brain Mapping, 45(7). 10.1002/hbm.26699

48. Taketomi, N., Michimae, H., Chang, Y.-T., & Emura, T. (2022). meta.shrinkage: An R package for meta-analyses for simultaneously estimating individual means. Algorithms, 15(1), 26. 10.3390/a15010026

49. Thulin, M. (2024). Modern statistics with R. CRC Press.

